# When the going gets tough, the tough get going: effect of extreme climate on an Antarctic seabird’s life history

**DOI:** 10.1101/791855

**Authors:** Stéphanie Jenouvrier, Lise Aubry, Silke van Daalen, Christophe Barbraud, Henri Weimerskirch, Hal Caswell

**Affiliations:** Biology Dept., MS-50, Woods Hole Oceanographic Institution, Woods Hole, MA 02543, USA; Centre d’Etudes Biologiques de Chizé, UMR 7372 CNRS / Univ La Rochelle-79360 Villiers en Bois, France; Fish, Wildlife and Conservation Biology Department, Colorado State University, Fort Collins, CO 80523-1474, USA; Institute for Biodiversity and Ecosystem Dynamics, University of Amsterdam, PO Box 94248, 1090 GE Amsterdam, The Netherlands

**Keywords:** unobserved individual heterogeneity, individual quality, frailty, fixed hetero-geneity, dynamic individual heterogeneity, stochasticity

## Abstract

Individuals differ in many ways. Most produce few offspring; a handful produce many. Some die early; others live to old age. It is tempting to attribute these differences in outcomes to differences in individual traits, and thus in the demographic rates experienced. However, there is more to individual variation than meets the eye of the biologist. Even among individuals sharing identical traits, life history outcomes will vary due to individual stochasticity, i.e., to chance. Quantifying the contributions of heterogeneity and chance is essential to understanding natural variability. Inter-individual differences vary across environmental conditions. Heterogeneity and stochasticity depend on environmental conditions. We show that favorable conditions increase the contributions of individual stochasticity, and reduce the contributions of heterogeneity, to variance in demographic outcomes in a seabird population. The opposite is true under poor conditions. This result has important consequence for understanding the ecology and evolution of life history strategies.

## 2 Introduction

There exist two sources of variation in life history outcomes: *individual heterogeneity* and *individual stochasticity*. Individual heterogeneity refers to differences among individuals in life history traits that in turn affect the vital rates to which they are subject. Heterogeneity may be fixed or dynamic (reviewed by [1, 2]). Fixed individual heterogeneity may be due to e.g. genetic variation, epigenetics, maternal effects, or permanent environmental effects. Dynamic individual heterogeneity may be due to changeable factors such as age, experience, health, or dynamic environmental effects [3]. Individual stochasticity is variability in demographic outcomes that is generated by random events in the life cycle (surviving or not, reproducing or not, etc.). Individuals will differ in their life trajectories and demographic outcomes, even if they are subject to identical vital rates, because of chance alone [4, 5, 6, 7, 3, 8, 9, 10].

The relative importance of these sources of variation in life history outcomes is critically important in improving our understanding of population dynamics and life history evolution, and is currently a topic of intense debate [11, 10, 12, 13, 2, 8, 14]. Both theoretical models and empirical studies have found that individual stochasticity contributes a large part of the variance in life expectancy and lifetime reproduction, regardless of the variation of life history traits and outcomes among individuals and regardless of life-history type (iteroparous or semelparous) [14, 15].

Some individual differences are readily observable and easily incorporated as state variables (e.g., age, stage, size) in demographic models. Other differences are latent or unobserved, variously referred to as frailty, latent heterogeneity, or individual quality (e.g., [16, 1, 17]). We refer to these as *unobserved heterogeneity*. Unobserved heterogeneity can obscure, or even reverse, patterns of survival and reproduction at the individual level, with consequences for population dynamics and our understanding of life history evolution [18, 19, 20].

Environmental conditions affect the expression of heterogeneity. Harsh conditions may remove frail individuals through physiological stress or increased resource competition. Harsh conditions may also inhibit breeding, so that only robust or high-quality individuals survive and breed successfully [21, 22, 23, 24]. Under favorable conditions, survival and breeding may be high regardless of heterogeneity. Extreme climatic events may act as important filters on the evolution of life histories [25, 26, 27].

The consequences of environmental conditions for life history outcomes such as life expectancy, lifetime reproduction, when unobserved heterogeneity is incorporated, are poorly known and challenging to calculate. Our goal here is to partition the variance of several life history outcomes between individual stochasticity and unobserved heterogeneity across different environmental conditions.

The Southern fulmar (*Fulmarus glacialoides*), an Antarctic seabird, is the first wild species for which the variance in life history outcomes was successfully partitioned into contributions from individual stochasticity and individual heterogeneity [19].

The Southern fulmar forages near the ice edge, which is an area of high productivity [28]. When sea ice concentrations are low, the distance from the colony to the ice edge is large and foraging trips are longer. As a result, adults bring less food to their chicks, which then fledge in poor body condition. This leads to reduced probabilities of breeding and breeding success and reductions in population growth rate. Following [28], we define three environmental conditions based on sea ice concentration: low, medium, and high (see Methods).

A capture-mark-recapture analysis including a latent heterogeneity variable revealed three groups with distinct sets of life-history traits and outcomes that occur together [19]. We describe there groups below. The groups were estimated as fixed (at birth) heterogeneity; models with dynamic heterogeneity were not identifiable. Under medium environmental conditions, heterogeneity explains only a small fraction of variance in life expectancy (5.9%) and lifetime reproduction (22%). Here, we characterize how environmental conditions affect unobserved heterogeneity in vital rates, life history outcomes, and growth rate in this population. We estimate the relative contributions of individual stochasticity and heterogeneity to variance in life history outcomes in the three environments.

## 3 Results

### (a) Fulmar life history: heterogeneity and stochasticity

We estimated a stage-classified matrix population model (with the states pre-breeders, successful breeders, failed breeders, and non-breeders); see Methods and [28, 19] for details. We identified heterogeneity groups within the population by accounting for unobserved heterogeneity in CMR analysis. Three such life-history *complexes* (sets of life-history characteristics that occur together through the lifetime of an individual [19]) exist, reminiscent of the gradient of life-history strategy observed among species (i.e., the slow-fast continuum; in birds: [29]; in mammals: [30, 31, 32, 33, 34]):

1. Complex 1 (14% of offspring at fledging) is a slow-paced life history where individuals tend to delay recruitment, recruit successfully, and extend their reproductive lifespan.
2. Complex 2 (67% of offspring at fledging) consists of individuals that are less likely to recruit, have high adult survival, and skip breeding often.
3. Complex 3 (19% of offspring at fledging) is a fast-paced life history where individuals recruit early and attempt to breed often, but have a short lifespan.

“Quality” is a multidimensional property in this model. Individuals in complexes 1 and 3 are “high-quality” individuals [1] because they produce, on average, more offspring over their lives than do individuals in complex 2. But complex 2 is made-up of individuals that experience the highest levels of adult survival. Accordingly, we will compare the life history complexes in terms of integrative demographic measures rather than abstract concepts like quality or frailty. Finally, under medium environmental conditions, stochasticity explains a large fraction of variance in life expectancy (94.1%) and lifetime reproduction (78%).

### (b) Environmental effects on life history heterogeneity

#### Vital rates, individual differences and environments

To compare vital rates across environmental conditions, we weighted the average vital rates by *π*, noted **E**_*π*_(*θ*). We found that for all three life history complexes, breeding and success probabilities all decrease when sea ice conditions are low, but adult survival remains unchanged (Fig. 1, Supplementary Table 2). For example, the average breeding probability of previous successful breeder (**E**_*π*_(*β*_2_)) is 0.85 for medium sea ice conditions but declines to 0.56 when sea ice conditions are low.

**Figure 1:**
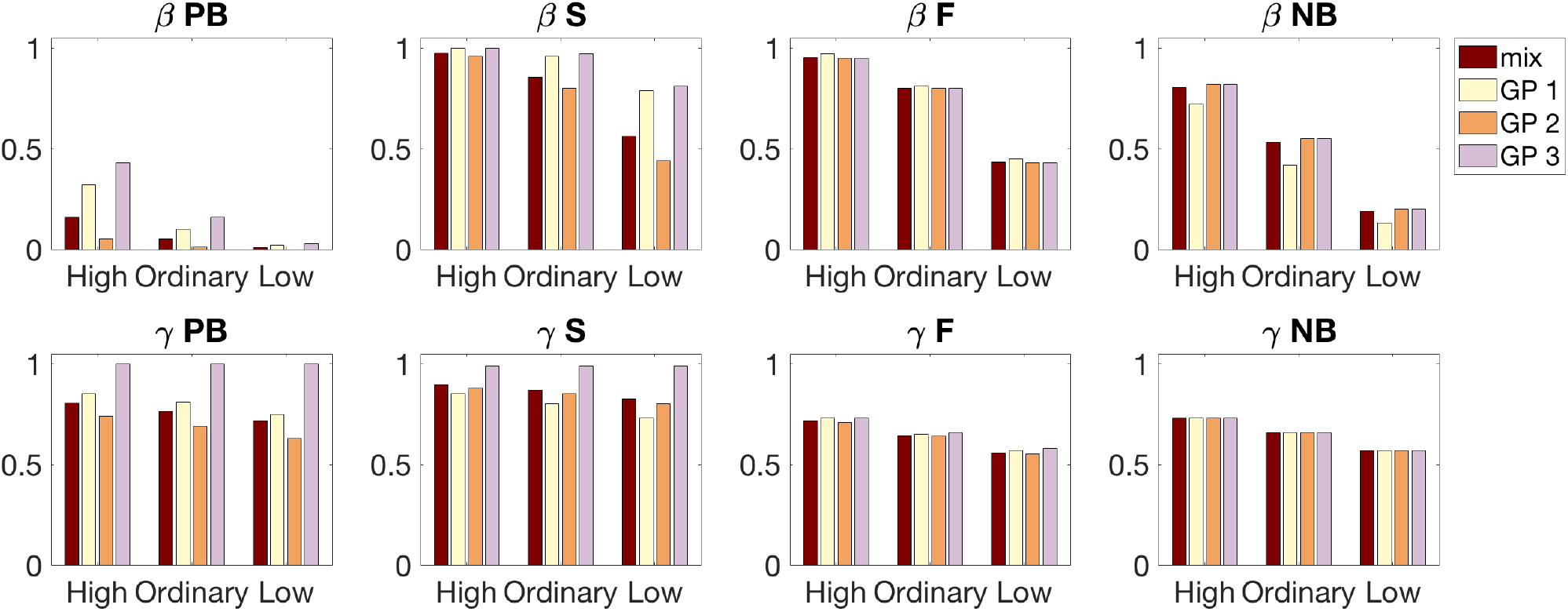
Vital rates of the southern fulmar for each reproductive state and sea ice conditions (SICs). Vital rates are averaged for environments characterized by: *high SICs* (1979, 1998, 2001), *low SICs* (1986, 1987, 2000) and *medium SICs* (all other years), as defined by [28]. Color bars refer to the 3 groups of unobserved heterogeneity (yellow: complex 1; orange: complex 2; and purple: complex 3), as well as the weighted average over the mixing distribution *π* = [0.14 0.67 0.19] (maroon). The panels are ordered by reproductive state at the previous breeding season (column 1: pre-breeders (PB); column 2: successful breeders (S); column 3: failed breeders (F); and column 4: non-breeders) and vital rates (first line: breeding probabilities *β*; and second line: success probabilities *γ*). Note that survival probabilities do not vary with time nor sea ice conditions, and thus are not shown here but in electronic supplementary material.

The impact of sea ice conditions on vital rates depends on the complex individuals belong to (Fig. 1). For example, the breeding probability of previously successful breeders (*β*_2_) decreases by *∼* 17% between medium and low sea ice conditions for complex 1 and 3 individuals, while it decreases by 45% for complex 2.

As a consequence, individual differences in vital rates depends on sea ice conditions. The coefficient of variation over the mixing distribution *π* measures these individual differences in vital rates, standardized by the mean, and weighted by *π* (Methods). We found that the difference in breeding and success probabilities among complexes increases when sea ice conditions are low (Fig. 2). Differences between complexes are more pronounced for the recruitment probabilities of pre-breeders, followed by the breeding probabilities of successful breeders, the breeding success of pre-breeders, and that of successful breeders.

**Figure 2:**
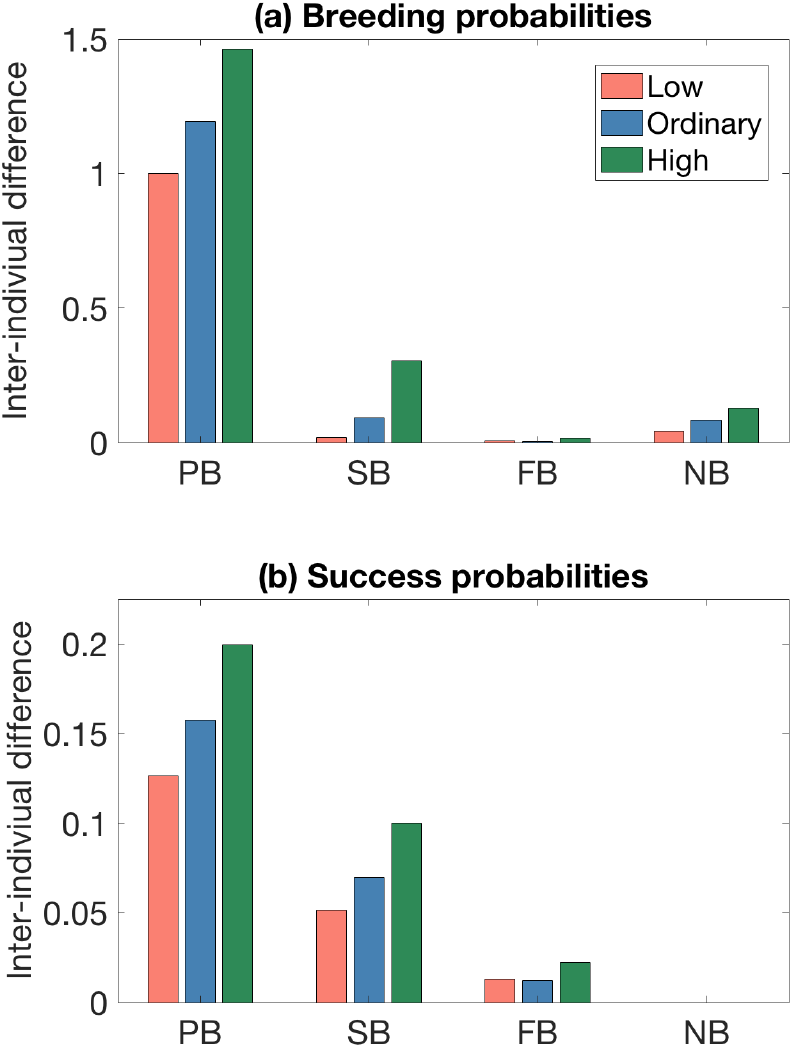
Individual differences in (a) breeding probabilities and (b) success probabilities across life history complexes for each set of sea ice conditions (SICs). Inter-individual differences are measured by the coefficient of variation over the mixing distribution. The x-axis indicates the reproductive state (see figure 1 for legends) and the bar colors refer to SICs (low: red, medium: blue and high: green).

#### Demographic outcomes, individual differences and environments

In each environment, the vital rates in each complex and the proportions of newborn individuals in each complex define means and variances (among individuals) in eventual demographic outcomes. We examine three integrative demographic outcomes: lifetime reproductive output, life expectancy, and population growth rate *λ*. These quantities are calculated for each environment, *as if* the population was living in such an environment permanently. This counterfactual calculation is typical of population projections being used to characterize the environment of a population by asking what would happen if that environment were maintained permanently [35].

We calculate demographic outcomes using the population projection matrix **A** for *λ* and the absorbing Markov chain implied by the matrix for calculating life expectancy and lifetime reproduction (see Methods). State transitions defined by the vital rates (Fig 1) and the time spent in each state (Fig. 4) interact to define life history outcomes (Fig. 3). For example, life expectancy varies across environmental conditions even if adult survival remains unchanged across sea ice conditions because adult survival differs among breeding states and the transitions among breeding states depend on sea ice conditions. The average life expectancy for complex 1 and 3 are larger for low sea ice conditions than for high sea ice conditions (Fig. 3) because they spend most of their life as pre-breeder (i.e. 85% and 79% of their lifetime, respectively; Fig. 4), a state that achieves higher levels of survival when compared to adults (Supplementary Table 2).

**Figure 3:**
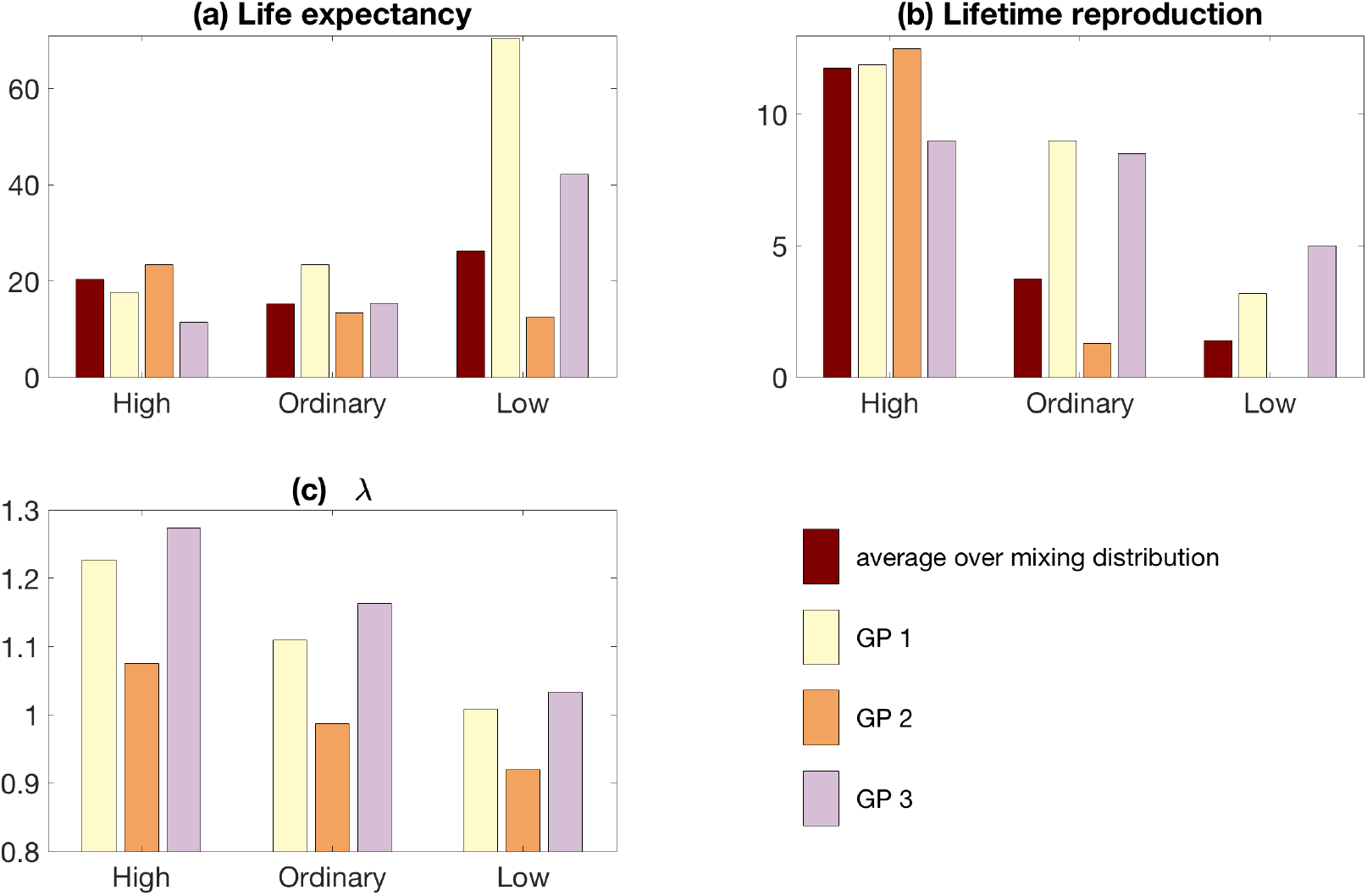
Demographic outcomes of Southern fulmar for each complex for each set of sea ice conditions. Color bars refer to the 3 groups of unobserved heterogeneity (yellow: complex 1; orange: complex 2; and purple: complex 3), as well as the weighted average over the mixing distribution *π* (maroon).

**Figure 4:**
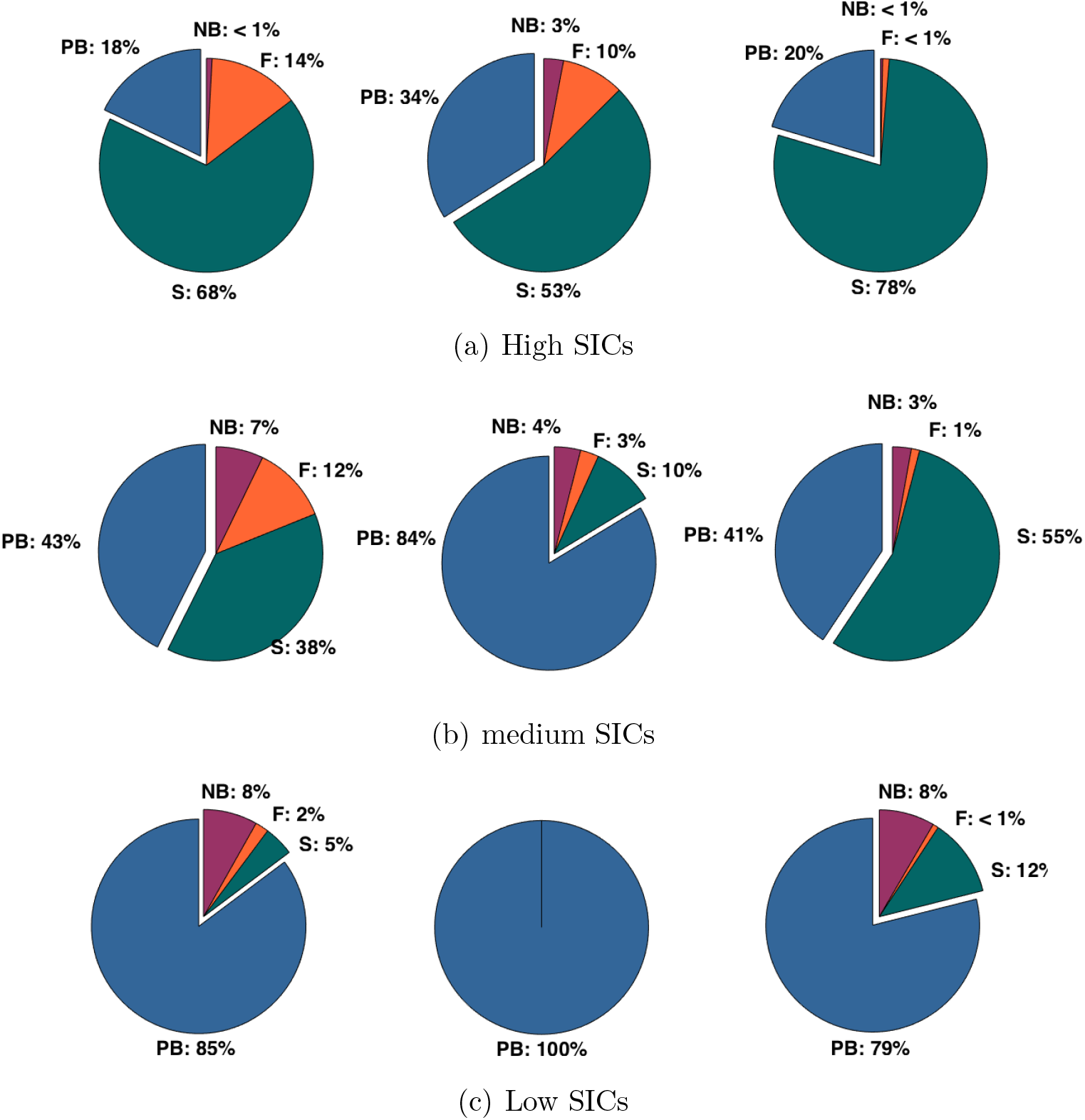
Percentages of time spent in each state for individuals in each complex for each set of sea ice conditions (a) high, (b) medium or (c) low. Complex 1 is shown by the left pie chart, while complex 3 is the right pie chart for each panel.

Overall, mean demographic outcomes across complex vary among sea ice conditions. The average life expectancy for the mixture of groups are larger for low sea ice conditions than for high sea ice conditions (red bars on Fig. 3). However, the average lifetime reproduction is larger when sea ice conditions are high and smaller when sea ice conditions are low.

The impact of sea ice conditions on life history outcomes depends on the complex individuals belong to. Individuals of complex 1 have the largest life expectancy, while individuals of complex 2 experience the shortest for an environment characterized by medium or low sea ice conditions (Fig. 3). In contrast, for high sea ice conditions, individuals of complex 2 achieve the highest life expectancy, while individuals of complex 3 experience the shortest. In such high sea ice conditions, individuals spend most of their life as adult breeders, and individuals of complex 2 have higher survival during adulthood than during the pre-breeding stage, while individuals that belong to life history complex 3 have the lowest adult survival.

For an environment characterized by medium or low sea ice conditions, individuals in complexes 1 and 3 produce, on average, more offspring over their lives than do individuals in group 2 (Fig. 3). When sea ice conditions are low, individuals in complex 2 are unlikely to recruit and their lifetime reproduction is null. However, for high sea ice conditions, individuals in complex 2 produce, on average, more offspring over their lives than do individuals in complexes 1 and 3, because they experience a longer lifespan.

Finally, we calculated the population growth rate *λ* to integrate all the rates into a measure that shows how successful a set of vital rates in one environment would be. Individuals of complex 3 have, on average, the highest *λ* regardless of environmental conditions (Fig 3).

### (c) Environmental effects on variance in demographic outcomes

The total variance in life history outcomes also varies across environmental conditions (Table 1). The total variance of life expectancy is larger when sea ice conditions are extremely low. On the other hand, the total variance of lifetime reproduction is much larger when sea ice conditions are extremely high.

**Table 1:** Life expectancy and lifetime reproduction for the southern fulmar in three environments characterized by high, medium or low sea ice conditions. Results of the variance partitioning of demographic outcomes between individual stochasticity and unobserved heterogeneity are also shown. % H is the percentage explained by unobserved heterogeneity.

|  | Total variance | Within-group | Between-group | % Heterogeneity |
| --- | --- | --- | --- | --- |
| HIGH |  |  |  |  |
| Life expectancy | 807.4 | 785.4 | 22 | 2.7 |
| Lifetime reproduction | 510.1 | 508.2 | 1.8 | 0.4 |
| medium |  |  |  |  |
| Life expectancy | 200.4 | 188.7 | 11.7 | 5.9 |
| Lifetime reproduction | 55.7 | 43.5 | 12.3 | 22 |
| LOW |  |  |  |  |
| Life expectancy | 1254.6 | 821.8 | 432.7 | 35.3 |
| Lifetime reproduction | 9.3 | 5.2 | 4.1 | 44.3 |

Both individual stochasticity and unobserved heterogeneity among individuals generate variability in life history outcomes [5, 14], in response to changes in both within-group (stochasticity) and between-group (heterogeneity) variances. For example, both the between-group and within-group variances of life expectancy increase when sea ice conditions are extreme. However, for an environment characterized by high sea ice conditions the increase in between-group variance is smaller than that in the within-group variance resulting in a smaller proportion of individual heterogeneity to the total variance.

In spite of these complex patterns, the proportion of variance in life expectancy and lifetime reproduction due to heterogeneity are smaller when sea ice conditions are high and larger when sea ice conditions are low. Indeed, partitioning the variance in life expectancy and lifetime reproduction reveals that only 2.7% of the variance in life expectancy and 0.4% of the variance in lifetime reproduction are due to individual heterogeneity when sea ice conditions are high, while 35.3% of the variance in life expectancy and 45.1% of the variance in lifetime reproduction are attributable to heterogeneity when sea ice conditions are low (Table 1).

## 4 Discussion

The impact of environmental conditions on unobserved heterogeneity in all fitness components of a species has virtually never been studied. Three different life-history *complexes* (sets of life-history traits that persist throughout the lifetime of an individual [19]) exist within a population of Southern fulmar. Here, we show that the differences in vital rates and demographic outcomes among complexes depend on the environmental conditions individuals experience. Importantly, differences across life history complexes are amplified when environmental conditions get extreme. Sea ice conditions did not only affect patterns of life history traits, but also the variance of life history outcomes and the relative proportion of individual unobserved heterogeneity to the total variance. These novel results advance the current debate on the relative importance heterogeneity (i.e. potentially adaptive) and stochasticity (i.e. enhances genetic drift) play in shaping potentially neutral vs. adaptive changes in life histories.

Our results indicate that extreme sea ice conditions affect vital rates, and the difference in vital rates among complexes depends on the environment fulmars live in (Fig. 1, 2). In years when sea ice conditions were low, fulmars traveled greater distances to forage and adults found less food to provision their chicks, ultimately affecting chick body condition and fledging success [28]. Fulmars feed mainly on krill (*Euphausia superba*) and other crustaceans, as well as on small fish (*Pleuragramma antarctica*) and squid [36]. During years with lower sea ice conditions the abundance of preys, such as krill, may be reduced [37, 38]. As a result, the breeding and success probabilities decline regardless of the complex individuals belong to, and differences in vital rates among life history complexes are larger when sea ice conditions are extremely low. Low sea ice conditions could intensify intra-competition for uneven resources and reveal differences among individuals of different “quality” [24, 39, 40, 1, 41].

Differences among life history complexes in vital rates vary among reproductive states. Individual differences in recruitment and success probabilities are larger for first-time breeders than experienced breeding adults (Fig. 2), probably because of pre-breeders’ limited experience with foraging in their ability to acquire, store, and conserve energy resources [42]. Individual differences in breeding and success probabilities are smaller for individuals which previously failed or skipped breeding when compared to individuals that previously succeeded. Raising an offspring successfully may impose an important energetic constraint on the probability of breeding (successfully) the following year and may intensify differences among complexes.

We also demonstrate that complexes differ in their demographic outcomes (life expectancy, lifetime reproduction, population growth rate *λ*), which further depend on the environmental conditions experienced (Fig 3). Complex 1 individuals (slow-paced life histories, with a delayed but high probability of recruitment and extended reproductive lifespan) have higher life expectancy and lifetime reproduction than any other complex when sea ice conditions are medium. Complex 2 individuals (low and delayed recruitment, skip breeding often, but with highest adult survival rate) have higher life expectancy and lifetime reproduction than any other complex when sea ice conditions are high, but a null lifetime reproduction and shortest life expectancy when sea ice conditions are low. Complex 3 individuals (fast-paced life histories), have higher lifetime reproduction than any other life history complex when sea ice conditions are low, and do achieve the lowest life expectancy in any environment.

Complex 3 individuals have the highest *λ* regardless of environmental conditions because they breed at younger ages than any other complex across all sea ice conditions. For Southern fulmars, recruitment probability is a key vital rate that has great potential in influencing population growth rate [28]. Extreme low sea ice conditions select for robust and high quality individuals (i.e complex 1 and 3 with *λ >* 1 Fig 3), because the competition for food resources increases. Thus “*when the going gets tough, only the tough get going*”. On the other hand, when sea ice conditions are extreme high, all individuals are more likely to access food ressources, hence survive and breed successfully, achieving a high fitness regardless of the complex they belong to.

Finally, our results indicate that the variance of life history outcomes depend on environmental conditions. The total variance in life expectancy are larger in extreme low sea ice conditions, while the total variance in lifetime reproduction is larger in extreme high sea ice conditions (Table 1). The total variance depends on the between and within groups variances, which are function of the variances and means of the life history outcomes of each group and the mixing distribution. The mean life expectancy of complex 1 and 3 are much larger in extreme low than high sea ice conditions, which may contribute substantially to the increasing total variance in life expectancy in low sea ice conditions. The mean lifetime reproduction is much larger in extreme high than low sea ice conditions for complex 2, which may contribute substantially to the increasing total variance in lifetime reproduction in high sea ice conditions.

Partitioning this total variance in demographic outcomes reveals that 35.3 and 45.1% of the variance in life expectancy and lifetime reproduction, respectively, is due to differences among complexes in a low sea ice condition, while only 5.9 and 22% of the variance is due to individual heterogeneity in a medium sea ice condition. This supports the hypothesis that more variability in life history outcomes is attributable to persistent intrinsic differences between individuals when competition intensifies for uneven resources [43, 24]. Indeed, differences across individuals in their ability to secure limited food resources may be exacerbated when sea ice conditions are low [28], leading to the observed increased contribution of individual heterogeneity to variance in life history outcomes.

In high sea ice conditions, 2.7 and 0.4% of the variance in life expectancy and lifetime reproduction, respectively, is due to differences among complexes. In high sea ice conditions, the foraging trips are shorter (sea-ice edge is closer to the colony) and food ressources likely more abundant [28]. Hence, more variability in life expectancy and lifetime reproduction is attributable to stochasticity under “favorable” conditions probably because all individuals survive and breed successfully regardless of the complex they belong to.

Our results based on a deterministic analysis in each environment place bounds on the degree to which individual heterogeneity can contribute to the variance of life history outcomes. However, individuals experience diverses environmental conditions during their lifetime. A stochastic model is required to partition the variance of life history outcomes between groups and environmental conditions. Furthermore, individuals may belongs to various complexes during their lifetime (i.e. dynamic individual heterogeneity). Unfortunately, a model to estimate transitions among unobservable states is not identifiable [19]. Further work entails exploring the consequences of such dynamic heterogeneity in a theoretical framework.

In conclusion, based on our findings in a long-lived vertebrate species, individual stochasticity makes a substantial contribution to variance in demographic outcomes when environmental conditions are favorable and medium, but individual heterogeneity contributes substantially to these outcomes when environmental conditions are poor. Because the strength of selection on fitness components often varies considerably from year to year in wild populations [44], we expect phenotypic selection on hidden traits to intensify when conditions are poor. These results advance the debate on how neutral versus potentially adaptive processes shape the variance of life history outcomes, and we further observe that the environmental context is key in molding the relative contribution of these process to the evolution of life histories. Finally, our findings support the hypothesis that both observed and unobserved differences across individuals can be tempered by environmental conditions, and ultimately define the diversity of life history strategies within a species.

## 5 Methods

### Definition of environmental conditions

Sea ice conditions affect the vital rates of Southern Fulmar [45], likely through their impact on food resources. In this population, individuals forage near the ice edge [28]. We use an index of sea ice condition which combines sea ice cover, and location of the sea ice edge (see [28] for more details on sea ice condition data and index calculation). We define low and high sea ice condition years as years with an index of sea ice conditions lower or higher than the 10th and 90th percentile of the sea ice condition distribution, respectively.

### Estimating unobserved heterogeneity in vital rates

To estimate both unobserved and observed sources of heterogeneity in vital rates, we use MultiState Mark-Recapture (MSMR) models with finite mixtures that account for imperfect detection [46, 47, 48] (Supplementary methods). Finite mixture MSMR models define a finite number of groups (hidden states) in the population and provide estimates for vital rates within each group. They also estimate the proportion of the sampled individuals falling into each heterogeneity group. We denote this distribution (the *mixing distribution*) ***π***. Mixture models have been intensively used in psychology, sociology, toxicology, and medicine to reveal the diversity of individual trajectories occurring within a population over time (see review in [48]). In ecology and evolution, few studies have used mixture models, and mainly for controlling for unobserved heterogeneity rather than trying to assess whether this heterogeneity influences within-population trajectories [48]. Here, we used these finite mixture MSMR models to reveal persistent life history complexes across contrasted environmental conditions.

We build on previous studies that identified 3 groups of individuals (i.e. life history complexes, [19]) based on unobserved heterogeneity in vital rates. We perform model selection to test for the effect of sea ice conditions on all vital rates of interest, once unobserved heterogeneity is accounted for. The best performing models selected (as measured by ∆*AIC*) comprised 90% of the overall AIC weight among the set of candidate models tested. The model-averaged vital rates are shown in figure 1.

### Coefficient of variation as a measure of differences among group

The coefficient of variation over the mixing distribution *π* is given by:

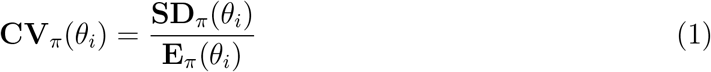

with *θ*_*i*_ a vector 3 *×* 1 of values for demographic rate *i* of the 3 groups of unobserved heterogeneity, for a specific environment. **E**_*π*_(*θ*_*i*_) is the mean across unobserved heterogeneity group weighted by the mixing distribution: **E**_*π*_(*θ*_*i*_) = *π*^T^*θ*_*i*_ and **SD**_*π*_(*θ*_*i*_) is the standard deviation calculated as:

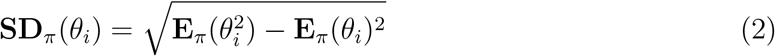

### Life history outcomes and their variance

We estimate life history outcomes (life expectancy, lifetime reproduction and age at first breeding) using absorbing finite-state Markov chains [49, 4] for the three environments characterized respectively by low, medium, and high sea ice conditions. We decompose the variance in life history outcomes into two components — individual heterogeneity and individual stochasticity. Associated equations are detailed for the Southern fulmar in an environment characterized by medium sea ice conditions in [19].

### Growth rate

We estimate individual fitness using a structured population model as the growth rate of a group of individuals with the same realized life history [50]. Thus we construct a population matrix for each complex (see supplementary material) and each set of environmental conditions, and calculate the deterministic growth rate as the maximum eigenvalue of this matrix [35].

## Data accessibility

Datasets will be available via Dryad following publication.

## Acknowledgments

We thank all the field workers who participated to the long-term study since 1964. We acknowledge Institute Paul Emile Victor (Programme IPEV 109), and Terres Australes et Antarctiques Françaises for for logistical and financial support in Terre Adélie. The Ethics Committee of IPEV and Comité de l’Environnement Polaire approved the field procedures. We acknowledge Dominique Besson and Karine Delord for fulmar data management.

## Authors’ contributions

SJ and HC conceived the ideas, designed methodology and obtained funding for the analyses; CB and HW collected the data and obtained funding for field-work; SJ and SVD performed preliminary analyses; SJ analyzed the data; SJ led the writing of the manuscript with LA and HC. All authors interpreted the data, contributed critically to the intellectual content and gave final approval for publication.

## Data accessibility

Data of this publication are archived at Dryad and available online at:

